# Large-scale assessment of olfactory preferences and learning in *Drosophila melanogaster*: behavioral and genetic measures

**DOI:** 10.1101/014357

**Authors:** Elisabetta Versace, Julia Reisenberger

## Abstract

In the Evolve and Resequence method (E&R), experimental evolution and genomics are combined to investigate evolutionary dynamics and the genotype-phenotype link. This approach requires many replicates with large population sizes, which imposes severe restrictions on the analysis of behavioral phenotypes. Aiming to use E&R for investigating the evolution of behavior in *Drosophila*, we have developed a simple and effective method to assess spontaneous olfactory preferences and learning in large samples of fruit flies using a T-maze. We tested this procedure on (a) a large wild-caught population and (b) 11 isofemale lines of *Drosophila melanogaster*. Compared to previous methods, this procedure reduces the environmental noise and allows for the analysis of large population samples. Consistent with previous results, we show that flies have a spontaneous preference for orange vs. apple odor. With our procedure wild-derived flies exhibit olfactory learning in the absence of previous laboratory selection. Furthermore, we find genetic differences in the olfactory learning with relatively high heritability. We propose this large-scale method as an effective tool for E&R and genome-wide association studies on olfactory preferences and learning.

## Introduction

Ongoing evolutionary dynamics and genotype-phenotype mapping can be studied during experimental evolution through subsequent phenotyping and genomic sampling [1,2]. This method is known as Evolve and Resequence (E&R) [3] and can be applied to entire populations by sequencing at the same times hundreds of individuals (Pool-seq, see Futschik and Schlötterer 2010) of the same population. Thanks to the advancements of high-throughput sequencing techniques, the E&R method has been used to track the changes in genomic composition not only across thousand of generations in bacteria [2] but also in eukaryotes with a fast life cycle, such as yeast [5] and fruit flies [6]. This approach provides a very promising opportunity to investigate the evolution of complex traits and their genetic architecture with a limited budget [7], thus paving the way to the analysis of the evolutionary dynamics of traits that cannot be inferred through fossil records, including complex behavioral phenotypes [8].

The first implementation of E&R on a complex behavior focused on phenotypic and genomic change in response to artificial selection for shorter/longer inter-pulse interval in male courtship song in *Drosophila melanogaster* [9]. In this study thousand of loci have been identified that responded to artificial selection and differed between populations selected for different behaviors. Similar outcomes, with thousand of alleles that significantly change in frequency between generations and treatments, have been found also for morphological or physiological traits [10,11], showing that the same methodological issues apply to behavioral and other traits. Despite the success in identifying some causative genes [e.g. 12,13], theoretical [6,14] and empirical evidence [11,15] has clarified that many of the significantly changed variants are in fact false positives derived by short or long-distance linkage disequilibrium. Another limit that E&R shares with other genome-wide approaches, is low statistical power in identifying unknown causative variants [e.g. 14,16].

Although haplotype-blocks can be used to study the dynamics of selected genomic regions during experimental evolution [15], an effective E&R study should primarily minimize the false positives rate and maximize statistical power. As shown in recent theoretical and simulation work [6,14,17,18], to reach this aim several issues have to be taken into account in the design of the experiment: (a) use a large starting population (possibly hundreds or thousands of individuals); (b) use a large population size; (c) use at least 5-10 replicate populations; (d) run the experiment for dozens of generations; (e) reduce linkage disequilibrium. In the light of this, during E&R researchers should phenotype and propagate thousands of individuals in multiple replicate populations for many generations [8]. In E&R studies a practical limitation is imposed by the stages of propagation and phenotyping: when flies have to be individually phenotyped and manipulated, the time and working load required can force a reduction of the census size. For this reason investigating behavioral traits, that often require a large effort in phenotyping, poses a methodological challenge.

In this work we focus on the development of a fast and reliable method for phenotyping and propagating fruit flies in E&R of olfactory behavior, in particular spontaneous olfactory preferences and olfactory learning. Fruit flies show complex behaviors, can be easily maintained at a large census size, have a fast generation cycle and low linkage disequilibrium [19], thus are a convenient model to investigate the evolutionary dynamics and genotype-phenotype map of behavioral traits.

Olfactory behavior (olfactory preferences and olfactory learning) is a good candidate for E&R investigation of both spontaneous and learned responses, because of its remarkable conservation between *Drosophila* and vertebrates [20,21] and the presence of standing variation for both olfactory preferences and olfactory learning [22–24]. Genetic variability in the experimental population is crucial to apply E&R to fruit flies, because within the time scale of feasible experiments (50 generations of selection take about two years to be completed) new mutations have little impact on evolutionary change.

Different preferences for specific odors and odor concentrations have been documented in *D. melanogaster* using T-mazes [e.g. 25,26,27]. Phenotypic variability in olfactory behavior is associated with polymorphisms that influence reactions to different compounds [28,29] but to date E&R has not been applied to odor preferences [see 30 for a similar idea]. Olfactory learning has been extensively studied at the behavioral, genetic and neurobiological level [20,31–33]. Wild and mutant flies have been tested in associative conditioning tasks, typically the association between an olfactory conditioned stimulus and an electric or mechanical aversive stimulus. In a first paradigm developed to measure olfactory learning, Quinn and colleagues [34] tested groups of about 40 flies. From a starting tube flies could approach a light source at the end of a second tube painted with odor A (or B). When flies entered the second tube an electric shock was delivered and could be associated with odor A (or B). After being returned to the starting tube, flies could enter a third tube containing odor B (or A), that was not associated with any electric shock. At test flies could choose to enter either a tube with odor A or one with odor B. A performance index compared the fraction of flies that avoided the unpaired odor and those which avoided the shock-paired odor. Not all flies explored the tubes, and their performance was affected also by phototaxis, thus this small-scale assay produced low learning scores. Tully and Quinn [35] modified the paradigm to test groups of about 100 flies in an apparatus in which odor A matched with pulses of electric shock was followed by odor B in the absence of electric shock. The odors were delivered by vacuum so that all flies were exposed to the odor-shock contingencies. After training, the flies were tested in a T-maze where could choose to approach either odor A or B. The learning scores for this procedure ranged between 0.7 and 0.9 [31] but the need of dedicated machines and hands-on operations on the flies limit the application of this method to large-scale long-term experiments.

To date the main method used in experimental evolution for enhanced learning [36,37] is the oviposition paradigm. This method is a medium/large-scale procedure based on the habit of flies to use the same medium for foraging and egg laying. The procedure starts exposing hundreds of free ranging flies to olfactory (or visual) stimuli associated to palatable or aversive media displaced in petri dishes located in a box (e.g. orange juice smell associated with palatable food, apple juice smell associated with aversive food). After exposure to olfactory (or visual) and associated gustatory stimuli flies are tested for their olfactory (or visual) preferences for olfactory stimuli previously associated or not associated with the aversive food. Compared to flies that do not remember the association, flies that remember the contingencies presented during the exposure phase are expected to lay more eggs in the substrate whose smell (or color) was never associated with aversive food. The proportion of eggs laid in the substrate associated with the palatable flavor is used as a proxy for learning.

To select for enhanced learning across generations, Mery & Kawecki [36] rinsed, moved to a neutral medium and propagated only the eggs laid in the medium previously associated with palatable food (alternatively Dunlap & Stephens [38] displaced eggs individually with a needle). As effect of this regime, in about 15 generations the proportion of flies which made the “correct” choice significantly increased. In spite of this, several aspects make the oviposition procedure less than ideal for E&R studies: (a) this method is prone to experimental noise, as shown by the lack of learning effects before selection; (b) only females are exposed to selection, because learning is measured using laid eggs, thus reducing the selective pressure to half of the propagated individuals and preventing to investigate behavioral domains different from oviposition and sex effects; (c) the oviposition paradigm imposes selection for fertility, egg laying during the few hours of the test and resistance to egg washing; (d) this paradigm does not control for the experience provided during the conditioning phases (many flies might not experience all stimuli before making a choice in the test phase); (e) extensive work to rinse/displace the eggs and propagate the flies is required, in turn reducing the number of experimental replicates that can be propagated.

Aiming to use E&R for investigating the evolution of behavior in fruit flies, we have developed a simple and effective method based on a T-maze to assess spontaneous olfactory preferences and learning in large samples (hundreds) of fruit flies of both sexes as well as in smaller samples (dozens of individuals). We find evidence for spontaneous and conditioned preferences for olfactory stimuli in a large population of *D. melanogaster* originally caught in South Africa and in a population of inbred lines originally caught in Portugal. Our procedure reduces the impact of undesired selective pressures and the effort in propagation and phenotyping. Furthermore this method is sensitive enough to detect learning, spontaneous preferences and heritability of these traits. We discuss the relevance of this T-maze based procedure for E&R and genome-wide association studies.

## Materials and methods

### Subjects

All experiments were run on isofemale lines of the same species, *Drosophila melanogaster*. Flies were maintained on standard cornmeal-soy flour-syrup-yeast medium, except during the experimental assays. Before the beginning of the experiments we kept all lines for at least two generations at 22°C in a constant 14:10 h light:dark cycle.

The population-wide experiments were ran on 670 lines derived from a natural population of *D. melanogaster* collected in Paarl (South Africa) in March 2012. In each trial we used a group of 250 adult flies (males and females 2 days old or older), randomly picked from the 670 isofemale lines of the South African population.

The individual-line experiments were ran on 11 inbred lines derived from a *D. melanogaster* population originally collected in Povoa de Varzim (Portugal) in July 2008. Before inbreeding, B101, B192 and B211 were maintained in the laboratory as isofemale lines. R1-R10 were derived from the same original population after being exposed to an experimental evolution procedure at different temperature regimes. The hot and cold temperature regimes are described in Orozco-terWengel et al. [10] and Tobler et al. [11]. Before inbreeding, R1-R5 entered the hot temperature treatment, whereas R6-R10 entered the cold temperature treatment. Inbred lines were generated through full-sib mating for 17 or more generations (B101: 17 generations; B192: 18 generations; B211: 19 generations; R1 and R3: 27 generations; R2: 29 generations; R5: 21 generations, R6 and R9: 20 generations; R7 and R10: 22 generations). For each line a virgin female and a randomly collected male were allowed to mate and from their offspring another virgin female and a random male were used to create the next generation. After inbreeding, these lines were kept as isofemale lines until the experimental assays. In each trial we used a group of 40 flies (males and females 2 days old or older) of the same line.

### Apparatus and stimuli

The T-maze (31 x 17.5 cm) used for the experimental assays (Fig. 1A) consisted of a starting chamber and a central chamber (12 x 8 x 1.5 cm) connected on each side to a food chamber. The starting chamber (9.5 x 2.5 cm) contained the flies at the beginning of each experimental phase. Food chambers (9.5 x 2.5 cm) were filled with experimental food. In each experimental phase flies begun the exploration of the apparatus from the starting chamber. The central chamber was connected to the food chambers with a funnel that prevents flies to re-enter the central chamber once they have approached the food. A similar trapping technique has been previously used for fruit flies.

**Fig. 1.**
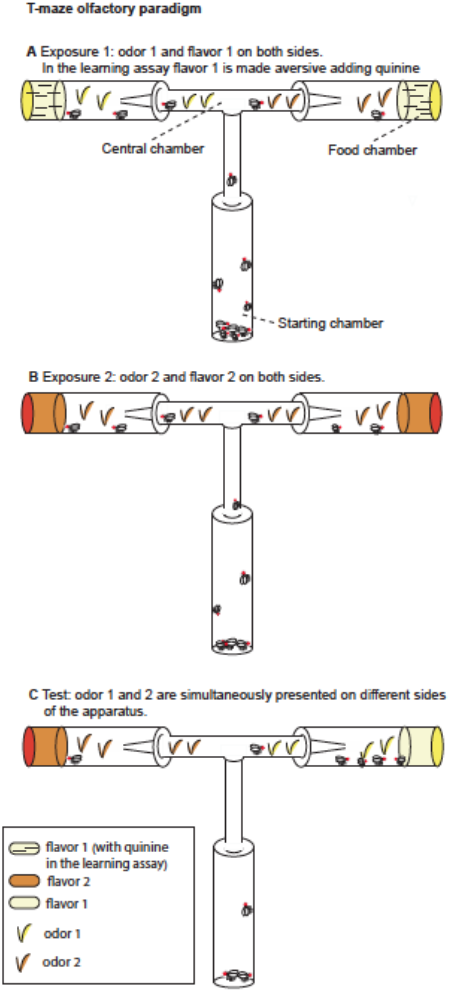
T-maze apparatus and experimental paradigm. **A** During Exposure 1 flies are presented with one odor (apple or orange) and the aversive food (apple or orange juice supplemented with quinine in the learning assay, without quinine in the spontaneous preference assay). **B** In Exposure 2 flies are presented the second odor (orange or apple) and the second flavor (orange or apple juice without quinine). **C** At Test both stimuli (orange and apple odor) are presented (without quinine) on different sides of the apparatus.

Experimental media prepared with juice fruit (either orange or apple juice from 100% concentrate) and agar (14 g/l). Aversive food was obtained adding 8 g/l of quinine to the experimental medium.

## Procedure

### Unconditioned olfactory preferences

We assessed unconditioned preferences for apple and orange odor by using the same procedure described for the learning assays (see below), with the only difference that no food supplemented with quinine was provided during the exposure phases. This similarity enables us to investigate the role of the conditioning procedure on spontaneous preferences.

### Olfactory learning assays

We used CO_2_ anesthesia to collect flies and starve them 15-16 hours before the beginning of the conditioning procedure. After starvation flies were moved to the starting chamber for Exposure 1. During Exposure 1, for two hours flies were exposed to the odor associated with the aversive flavor and the aversive flavor (e.g. orange odor and orange juice supplemented with quinine). Flies who entered the food chambers during Exposure 1 were moved to the starting chamber for Exposure 2. During Exposure 2, for two hours flies were exposed to the odor associated with the palatable flavor and the palatable flavor (e.g. apple odor and apple juice). Flies who entered the food chambers during Exposure 2 were starved for four hours prior to the Test.

In half trials we conditioned flies on apple odor associated with aversive food and orange odor associated with palatable food (A-/O), in half trials we conditioned flies on orange odor associated with aversive food and apple odor associated with palatable food (O-/A). It has been shown that in *D. melanogaster* appetitive long-term memory occurs after single-cycle training also in the absence of fasting [39], while aversive long-term memory by single-cycle training requires previous fasting [40]. For this reason the exposure to the aversive stimulus was always conducted immediately after fasting (Exposure 1).

Flies began the Test from the starting chamber. Differently from the exposure phases, during the Test the odor associated with the aversive flavor and the odor associated with the palatable flavor were presented simultaneously, each on a different food chamber (Fig. 1C). We alternated the right/left side in which the two odors were presented. No food was supplemented with quinine during this phase. We counted flies that chose to enter either the orange odor side or the apple odor side.

### Data analysis

In the test for spontaneous preferences between the orange and apple odor we compared the proportion of flies that across 28 trials chose the orange odor vs. the random choices level using a t-test single sample against the random choices proportion of 0.5. Beforehand we controlled for deviations from the normal distribution of the data using the Shapiro-Wilk normality test. To check for any effect of the order of presentation (O/A: Orange followed by Apple odor, A/O Apple followed by Orange odor) we calculated the order score *o* as the difference in the proportion of flies that in each trial chose orange odor after being exposed to O/A *vs*. A/O:

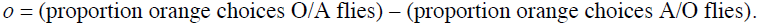

An order score significantly different from zero was expected if the order of presentation of the two odors/flavors had an effect.

Similarly, in the test for conditioned preferences between the orange and apple odor we compared the proportion of flies that across 28 trials chose the orange odor vs. the chance level using a t-test single sample against the random choices proportion of 0.5. Beforehand we controlled for deviations from the normal distribution of the data using the Shapiro-Wilk normality test.

To obtain a measure of learning (learning score, *l*) we calculated the difference in the proportion of flies that in each trial chose orange odor after being conditioned on apple flavor *vs*. orange flavor:

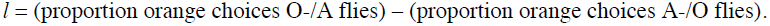

A learning score significantly different from zero was expected if the conditioning had an effect.

To investigate the heritable component of learning we repeatedly tested the same inbred lines (*n*=10) and derived the intraclass correlation t, which is an estimate of the genetic heritability (*h*^*2*^) of learning for the tested population, using the variance between (Vb) and within (Vw) inbred lines [41]:

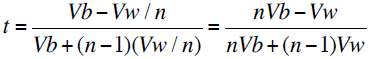

As discussed by David et al. [42] the intraclass correlation can be used as a proxy for heritability [see also 41,43].

## Results

### Population experiments: spontaneous preferences and learning assays

To evaluate the sensitivity of our method in detecting learning and spontaneous preferences in large groups of naturally derived fruit flies, we investigated the differences in olfactory learning and spontaneous preferences in a large South African *D. melanogaster* population. In each trial we tested 250 flies of both sexes.

In the spontaneous preference experiment, across 28 test trials the overall population showed a marginally significant preference for the orange odor (t_27_=1.953, p=0.06). This preference is consistent with the preference for citrus previously documented in *D. melanogaster* [36 but see Betti et al. 2014, e.g. 44], that is likely a behavioral strategy against the infection of parasitic wasps [44].

Before testing flies, we exposed them to both odors/flavors: in half trials flies were exposed first to orange then to apple (O/A), in half trials first to apple then to orange (A/O). We have derived the order effect score *o* to investigate the effect of the order in which the orange/apple stimuli had been presented. We observed a significant order effect score (t_13_=3.09, p=0.009; Fig. 2B), indicating that A/O flies (flies first exposed to Apple, then to Orange) had a significantly higher preference for orange odor than O/A flies (flies first exposed to Orange, then to Apple). A post-hoc t-test on A/O and O/A flies vs. the chance level (0.5) revealed that only A/O flies had a significant preference for the orange odor: for A/O flies t_13_= 3.18, p= 0.007; for O/A flies t_13_= -0.242, p=0.81.

**Figure 2.**
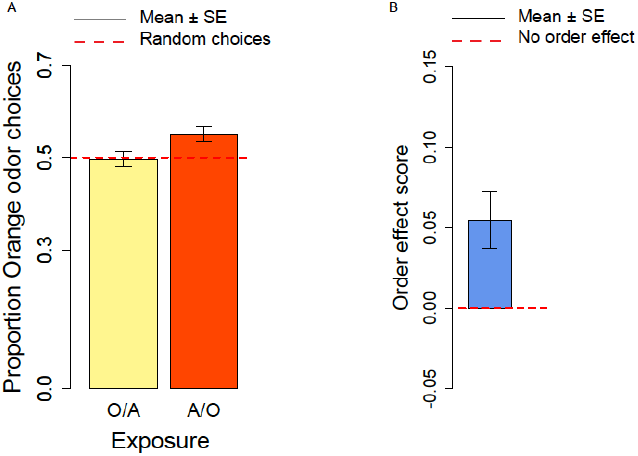
Spontaneous orange choices and order effect score. **A** Proportion of orange choices of flies exposed to Orange as first, Apple as second stimulus (O/A), and to Apple as first, Orange as second stimulus (A/O). **B** Order effect score (difference in orange odor choices between flies exposed to A/O and O/A).

In the conditioning experiment the overall population showed a preference for the orange odor (mean=0.59, t_57_=3.95, p<0.001). Flies that previously experienced Apple as aversive/Orange as palatable (A-/O) were more likely to choose orange than flies exposed to the opposite contingency (O-/A) (t_56_=2.24, p=0.029; Fig. 3A). The population showed a significant learning score (t_28_=2.88, p=0.007; Fig. 3B).

**Figure 3.**
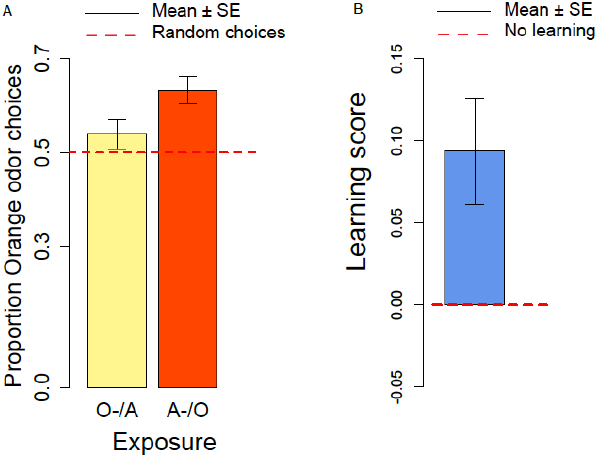
Orange choices after conditioning and learning score. **A** Proportion of orange choices of flies conditioned with Orange aversive/Apple palatable (O-/A, yellow) and Apple aversive/Orange palatable (A-/O, orange). **B** Learning score: difference in the proportion of orange odor choices between flies conditioned A-/O (Apple aversive, Orange palatable) and O-/A (Orange aversive, Apple palatable).

We also checked for differences in the proportion of orange choices after the conditioning procedure and after the spontaneous preference exposures. Overall, after conditioning flies had a stronger preference for orange odor than in the absence of conditioning (t_81_=2.50, p=0.014). These results suggest that exposure to aversive stimuli can influence the preferences of flies towards specific odors/flavors.

When comparing samples that had the same order of presentation of apple and orange odor in the conditioning and spontaneous preferences procedure, we observed a significant difference for the A/O but not for the O/A presentation (A-/O: t_40_=2.49, p=0.017; O-/A: t_39_=1.24, p=0.22). These results indicate that the conditioning procedure is more effective for the A-/O exposure than for the O-/A exposure.

### Inbred lines: spontaneous preferences and learning assays

To study olfactory preferences and learning in 11 inbred lines of *D. melanogaster* derived from a population collected in Portugal we used the same procedure adopted for the large population using 40 flies from the same isofemale line in each trial. Line R6 consistently did not enter the food chambers in both experiments, so we excluded this line and run the analyses in the ten remaining lines.

In the spontaneous preference assay the overall distribution of the orange odor choices was significantly different from the normal distribution (Shapiro-Wilk normality test: W=0.98, p=0.03) and we analyzed the data using non-parametric tests (Wilcoxon signed-rank test and Kruskal-Wallis test). Overall, the group of ten responsive lines showed a spontaneous preference for the orange odor (mean=0.56; V=7862, p<0.001; Fig. 4A). We did not observe significant differences across lines (Kruskal-Wallis Chi squared_9_=14.14, p=0.12; see Fig. 4B) and in the effect of the order of presentation (A/O *vs*. O/A) (Kruskal-Wallis Chi squared_9_= 1.48, p=0.22).

**Figure 4.**
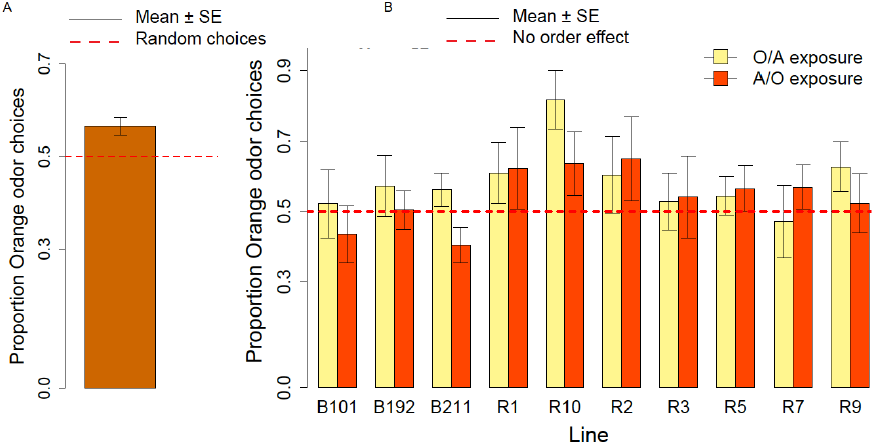
Spontaneous orange odor choices for a population of ten inbred lines and individual lines. **A** Overall proportion of spontaneous orange odor choices for a population of ten inbred lines. **B** Orange odor choices after exposure to Orange first Apple second (O/A) and to Apple first, Orange second (A/O) in each of ten inbred lines.

We calculated the order effect score – (proportion of orange odor choices after A/O exposure) – proportion of orange odor choices after O/A exposure – for the overall sample of ten lines tested and found no significant effect (V=1300.5, p=0.23).

In the learning assay, the overall distribution of the orange odor choices was significantly different from the normal distribution (Shapiro-Wilk normality test: W=0.98, p=0.005) and we analyzed the data using non parametric tests (Wilcoxon signed-rank test and Kruskal-Wallis test). Overall, after the conditioning procedure the ten responsive lines showed a preference for orange odor (mean=0.59, Wilcoxon signed-rank test, V=8630, p=9.159 x^10-6^), Fig. 5A. No significant difference in the overall choices was observed between the spontaneous preference and the learning assay (W=13655.5, p=0.30). Differently from the spontaneous preference assay though. significant differences in the proportion of orange choices were apparent between the two conditioning treatments (A-/O vs. O-/A: Kruskal-Wallis Chi square: 34.93, p=3.424 x^10-9^) (Fig. %A), and we documented a significant learning effect (Fig. 5B). We also detected significant differences in the proportion of orange choices between lines (Kruskal-Wallis Chi square: 23.45, p=0.005).

**Figure 5.**
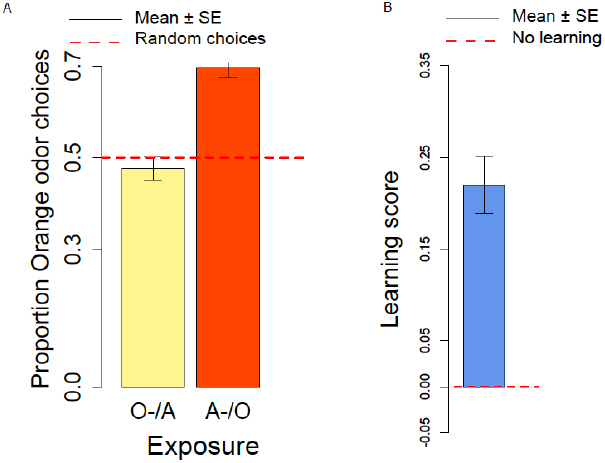
Conditioned orange odor choices for a population of ten inbred lines and learning score. **A** Proportion of orange odor choices for flies conditioned with Orange aversive/Apple palatable (O-/A) and Apple aversive/Orange palatable (A-/O) in the overall sample of ten inbred lines. **B** Learning score: difference in the proportion of orange odor choices between flies conditioned on A-/O (Apple aversive, Orange palatable) and O-/A (Orange aversive, Apple palatable).

We ran post-hoc tests to evaluate the performances of each line in the A-/O and O-/A conditioning procedure. For each line we measured the proportion of orange choices after conditioning with A-/O and O-/A (fig. 6A and B). After using the Bonferroni-Holmes correction for multiple comparisons, we found that in the A-/O procedure three lines (R10, R7, R9) had a preference significant at 5% level for orange and four lines (R1, R2, R3, R5) had a preference significant at 10% level, whereas in the O-/A procedure only R5 had a preference significant at 10% level. These results indicate that most of the tested inbred lines are able to discriminate between apple and orange odor.

**Figure 6.**
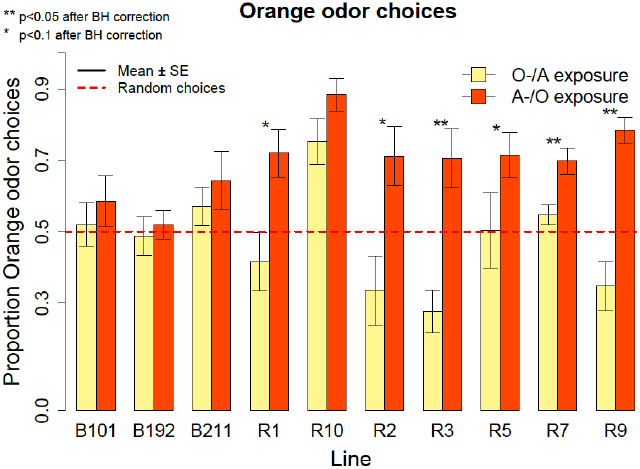
Proportion of orange odor choices for each tested line conditioned with Orange aversive/Apple palatable (O-/A exposure) and Apple aversive/Orange palatable (A-/O exposure).

We calculated the learning score – (proportion of orange odor choices after A-/O exposure – proportion of orange odor choices after O-/A exposure) – for the overall population and found a significant effect of learning (mean=0.22, V=2752.5, p<0.001). Since there was a significant effect of Line in the learning score (Kruskal-Wallis Chi square=22.15, p=0.008) we ran also post-hoc tests on each line (fig. 6). All lines had an higher proportion of orange choices after being conditioned with Orange as palatable stimulus (and all lines increased the proportion of orange odor choice after conditioning compared to the spontaneous choice assay, see Fig. 4B). Using the Bonferroni-Holmes correction for multiple comparisons we found that three lines (R3, R7, R9) showed a learning score significant at 5% level and three lines (R1, R2 and R5) that were significant at 10% level. These results suggest that most of the tested lines are able to learn through our conditioning procedure.

### Inbred lines: heritability of olfactory behavior

We derived an estimate of the genetic heritability of olfactory preferences and olfactory learning using the variance between (Vb) and the variance within (Vw) lines and calculating the intraclass correlation *t* as a proxy for heritability in inbred lines [41,42].

In the olfactory preferences, the variability between lines (Vb=0.09) was larger than the variability within lines (Vw=0.06) and the intraclass correlation is *t*=0.6. The same pattern holds true for olfactory learning: the variability between lines (Vb=0.017) is much higher than the variability within lines (Vw=0.004), thus leading to *t*=0.80. The high intraclass correlations show a moderate to high heritability of olfactory preferences and learning and suggest that our method is suitable to investigate these traits.

## Discussion

Historically the evolutionary dynamics of behavioral traits have been particularly hard to catch. This is not only due to lack of fossil record as a tool to help reconstructing evolutionary change but also to limits in investigating organisms with a complex behavior for hundreds or thousands of generations, as can be done with yeast [e.g. 45] and bacteria [2]. *Drosophila* is a model system which shows either complex behavior and a life cycle fast enough for being studied in experimental evolution. For instance, in few generations of targeted selection it has been possible to obtain a significant increase in learning in a wild-derived population of *Drosophila melanogaster* [36], or to change its responsiveness to specific (odor-flavor or color-flavor) associations [37]. These behavioral findings have not been accompanied by a correspondent genomic investigation, partly due to the costs and difficulties associated until recently to this enterprise. The recent development of high-throughput sequencing technologies, together with advancements in statistical and bioinformatics tools, has changed this scenario. In particular, using the Evolve and Resequence method [3] entire populations can be propagated and investigated for genomic changes at subsequent time points by sequencing collectively hundreds of individuals (a method known as Pool-seq, see Futschik and Schlötterer [4]). This approach paves the way to the analysis of complex behavioral phenotypes such as olfactory preferences and learning in *Drosophila* [8].

Empirical [9–11,15] and theoretical studies [6,14] have shown the current limits of E&R in terms of false positive and false negative errors. Different strategies have been suggested to reduce the error rate and increase the efficiency of this method in investigating evolutionary dynamics and the genotype-phenotype link [6,14,17], including propagate and phenotype large samples of several replicate populations for multiple generations. The oviposition method [36] is the experimental paradigm currently used for experimental evolution of learning in fruit flies [36–38]. This procedure though is not optimal for E&R due to several drawbacks: the effort required for propagation (e.g. eggs have to be rinsed and/or individually displaced on culture media), the fact that selection is imposed only on half of the propagated subjects (females) and males or male/female interactions cannot be investigated, the presence of selection for fast egg laying and survival to egg washing. With the aim of overcoming these limitations we have established a method based on subsequent exposure and test in a T-maze used to assess olfactory preferences and learning in large (hundreds or even thousands of individuals) and small (dozens) samples of fruit flies. This simple procedure can impose selection on both sexes and does not entail selection for fast egg laying and egg washing to propagate flies.

We have used the T-maze procedure to investigate spontaneous and learned olfactory choices in a large population of *D. melanogaster* originally caught in South Africa and in 11 inbred lines of another population of *D. melanogaster* originally caught in Portugal. Overall both populations show a spontaneous preference for orange vs. apple odor. The preference for citrus media had been previously documented in *D. melanogaster* [36, but see 43,44]. We find a significant effect of learning in both populations. The presence of olfactory learning in the absence of selection for this trait shows the sensitivity of our method.

Small population size and inbreeding negatively affect the resolution of genomic scans [6,14,15], thus limiting the power of E&R and genome-wide association studies. This limitation could be overcome using the T-maze procedure on large samples, since no individual handling of eggs is necessary in this method.

Moreover, repeatedly testing inbred lines we have detected genetic differences in olfactory behavior between lines. Differences between wild-derived inbred lines have been previously documented [46,47]. We have also calculated the intraclass correlation *t* [41,42] as an estimate of heritability, showing a medium/high heritability for the investigated traits.

We have showed that our method is suitable to be used with large samples to investigate the evolution of spontaneous preferences and learning performances in large groups of fruit flies with limited effort. On this basis we suggest to use of T-mazes in large-scale experiments as a tool for E&R and genome-wide association studies on olfactory preferences and learning and for other traits. Our experimental paradigm can be easily adapted to assess visual behaviors of fruit flies by changing the color/texture of the central and food chambers, and navigation performance by increasing the number of food chambers, as well the role of social information transmitted [see for instance 48,49] by one or both sexes. Varying the delay between conditioning and test it is possible to investigate the duration of memory. We suggest the use of large-scale T-mazes to widen the investigation of behavioral traits and cognitive abilities in invertebrates at the behavioral and genomic level.

## Acknowledgements

We are grateful to Christian Schlötterer for his support through all stages of the project. Special thanks to the members of the Institut für Populationsgenetik at Vetmeduni and the Vienna Graduate School of Population Genetics for helpful discussions throughout the project. We wish to thank Lino Ometto for fruitful discussions on the genetic analyses. The project has been funded through a grant of the Austrian Science Fund (FWF, P22725) awarded to C. Schlötterer.

